# A plausible mechanism for longitudinal lock-in of the plant cortical microtubule array after light-induced reorientation

**DOI:** 10.1101/2020.11.01.362285

**Authors:** Marco Saltini, Bela M. Mulder

## Abstract

The light-induced reorientation of the cortical microtubule array in dark-grown *A. thaliana* hypocotyl cells is a striking example of the dynamical plasticity of the microtubule cytoskeleton. A consensus model, based on *katanin*-mediated severing at microtubule crossovers, has been developed that successfully describes the onset of the observed switch between a transverse and longitudinal array orientation. However, we currently lack an understanding of why the newly-populated longitudinal array direction remains stable for longer times and re-equilibration effects would tend to drive the system back to a mixed orientation state. Using both simulations and analytical calculations, we show that the assumption of a small orientation-dependent shift in microtubule dynamics is sufficient to explain the long term lock-in of the longitudinal array orientation. Furthermore, we show that the natural alternative hypothesis that there is a selective advantage in severing longitudinal microtubules, is neither necessary nor sufficient to achieve cortical array reorientation, but is able to accelerate this process significantly.

## I. INTRODUCTION

The cortical microtubule array (hereafter CA) is a highly ordered structure occurring in cells of higher plants, which plays a key role in the growth-driven morphogenesis of cells and, therefore, also of the organism as a whole. It is known that the function of the CA is intimately linked to its spatial organization which, in turn, is linked to its dynamic properties [1]. In contrast to animal cells, the CA is built and reorganized without benefit of a microtubule organizing centre (centrosome). Instead, new microtubules in the CA are generated in two distinct ways: from nucleation templates - i.e., *γ*-tubulin complexes, mostly located on the lattice of already existing microtubules [2, 3], or from severing events of already-existing microtubules. The latter mode of generation of new microtubules is mediated by the severing protein *katanin* that localizes at microtubule crossovers and preferentially severs the newer one, i.e., the overlying one [4]. It plays a crucial role in the reorientation of the cortical array as a response to blue light. Typically, the direction of the cortical microtubule array of growing plant cells is transverse to the long axis of the cell. However, upon exposure to blue light, in dark-grown hypocotyl cells of *Arabidopsis thaliana*, the initially transverse array undergoes a striking reorientation to a direction longitudinal to the long axis of the cell. Since the initial array is transverse to the long axis of the cell, newer microtubules at crossovers are most likely longitudinal and, hence, the occurrence of multiple severing events quickly generates a new exponentially growing population of longitudinal microtubules [4].

Experimental and computational studies have identified the crucial role of the stability of the dynamic microtubule ends in enabling CA reorientation [5, 6]. Furthermore, a recent theoretical study has shown that the combination of preferential severing and a high probability of stabilization-after-severing for the newly-created microtubule plus-ends is a necessary ingredient for the reorientation process to start [7]. However, these studies focused on the first phase of the reorientation process, when the initial transverse array can still be seen as a constant background and free tubulin to build new microtubules is likely available in abundance. While these assumptions are realistic at the start of the reorientation process, at a later phase the amount of free tubulin is bound to become scarce as the number of growing microtubules increases. As transverse microtubules are also dynamic, this scarcity of tubulin would imply their gradual depolymerization, as they are outcompeted by the large population of growing longitudinal microtubules. This in turn, however, decreases the opportunity to create new crossovers and, therefore, to create new longitudinal microtubules from severing events. This suggests that while the initial asymmetry in the number of preferential severing of microtubules in the longitudinal direction is a sufficient ingredient to yield an initial asymmetry between the two populations of differently oriented microtubules, over time this bias is expected to fade. New transverse microtubules will start to be nucleated, and their crossovers with preexisting longitudinal microtubules can now serve to generate more transverse ones, through the same severing mechanism. In fact, all things being equal, one would expect the system to evolve to a novel steady state with an equal number of transverse and longitudinal microtubules. This clearly contrasts with the experimental findings of an ultimately stable longitudinal array, and raises the question of the mechanism by which this switch in orientations can be maintained.

Here, we explore the hypothesis that the ultimate asymmetry between differently oriented microtubules could be a consequence of small differences in their dynamic behaviour. The idea that the dynamics of microtubules can be influenced by cell geometry has received a lot of attention since the seminal work of Hamant et al. [8] (for a review see [9]) which provided the first evidence that microtubule would in fact prefer to align in the direction of maximal stress in the cell wall. In the most common cylindrical plant cell geometry this direction of maximal stress follows the direction of strongest curvature, which thus provides a possible explanation for the ubiquitous transversality of the CA in this cell geometry. Although the severing protein katanin is implicated in mediating the interaction between wall stress and microtubule stabilisation [10], the precise mechanism by which this occurred has not been elucidated to date. Computational models of cortical microtubule dynamics on closed surfaces with different geometries have also shown that slight directional cues, caused, e.g., by catastrophes induced at sharp cell edges or relative stabilization on specific cell faces, are sufficient to select a single preferential direction of ordering of the CA as a whole [11, 12]. On the basis of these considerations, we posit that a mechanism by which microtubule dynamics is sensitive to orientation with respect to a cell axis is possible, and that such a bias can impact the global organisation of the CA. Specifically, we will assume that microtubules in the longitudinal direction will grow slightly faster than ones in the transverse direction, but, as we show, an analogous decrease of catastrophe rate in the longitudinal direction will serve the same purpose. We implement our hypothesis in a stylized stochastic model of dynamic microtubules that can only occur in two distinct orientations - i.e., transverse and longitudinal. These two populations of microtubules compete for the same pool of available tubulin dimers as their building material. Besides the indirect interaction through the available tubulin pool, the two populations of microtubules directly interact through severing events at crossovers. Both simulations and additional analytical calculations show that the small difference in the dynamic parameters between the two populations can explain the experimentally observed lock-in of the CA to the longitudinal direction after reorientation from an initially transverse state. We also test an alternative hypothesis that the preferential severing of longitudinal microtubules is specifically required for the reorientation to occur. While we show that the latter mechanism is neither a necessary nor a sufficient ingredient for the reorientation to occur, we do find that it can significantly increase the speed of the reorientation.

## II. METHODS

To test our hypothesis that the full and subsequently maintained reorientation of the CA (see Figure 1A) is caused by a small asymmetry in the dynamics of differently oriented microtubules, we introduce a stochastic model for microtubules undergoing dynamic instability. We focus on the generation of new microtubules through nucleation and severing. A full mathematical description of the system of differential equations controlling the dynamics of individual microtubules and the steady-state solution for microtubule length distribution can be found in the Supplementary Information IX A.

**Figure 1.**
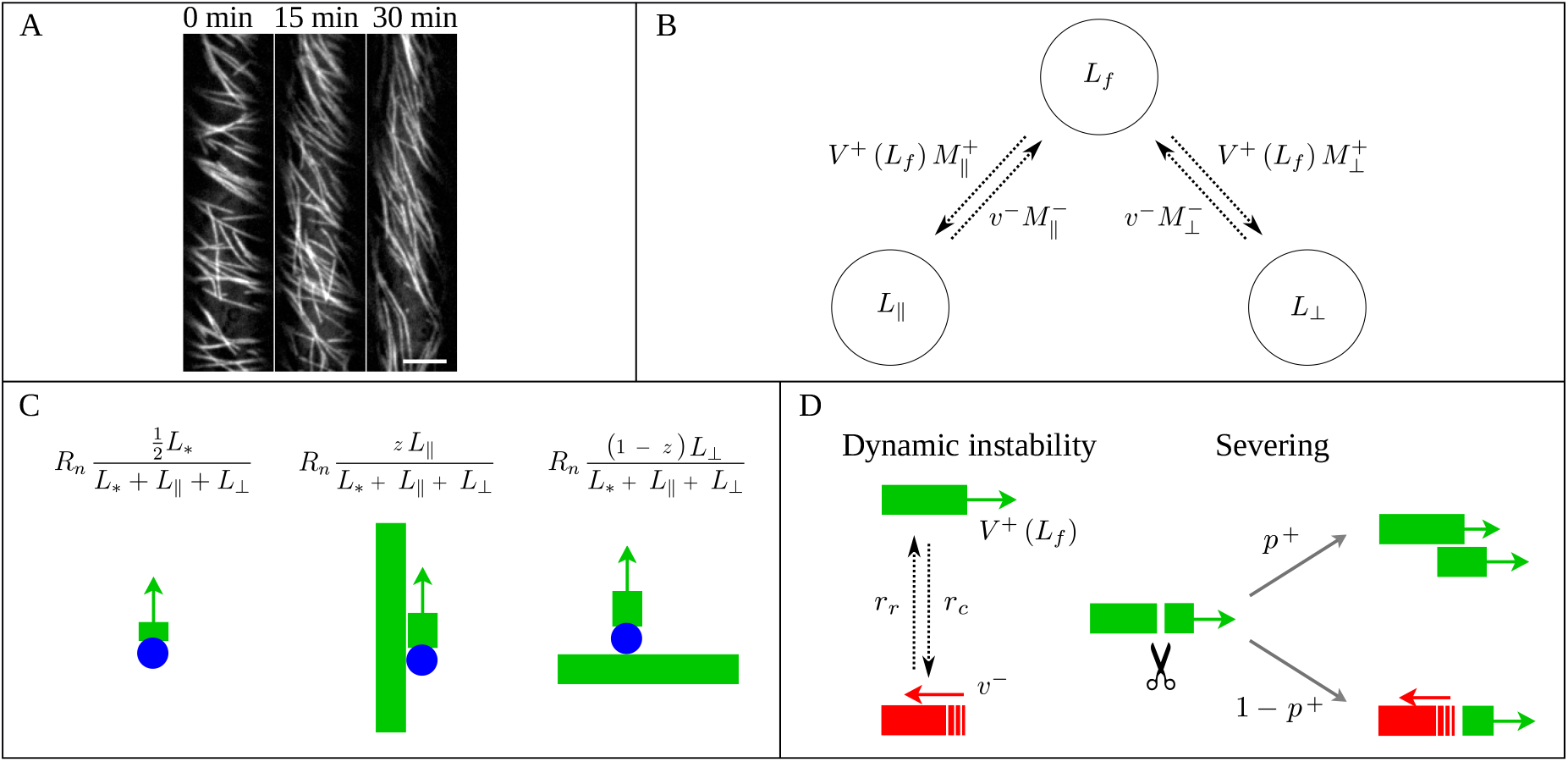
(A) Microtubule reorientation of dark-grown hypocotyl cells at 0, 15, and 30 min after induction of reorientation by blue light. Scale bar, 5 *µ*m. (B) Overall dynamics of the two microtubule populations and tubulin redistribution in the system. Free tubulin is recruited by microtubules of the two dynamic populations with a speed proportional to the total number of growing microtubules, while it returns to the free pool with a speed proportional to the total number of shrinking microtubules. The label *±* stands for growing/shrinking, respectively. (C) Nucleation rates for new longitudinal microtubules. From left to right, new microtubules can be nucleated through dispersed nucleation, microtubule-based nucleation parallel to the mother longitudinal microtubule, or microtubule-based nucleation orthogonal to the mother transverse microtubule. Blue circles represent the *γ*-tubulin complex. (D) Dynamics of an individual microtubule. Microtubules undergo dynamic instability and are severed with rate proportional to their length. Newly-created plus end after severing enters either the growing state with probability *p*^+^, or the shrinking state with probability 1 − *p*^+^.

### The model

The model, based on the Dogterom-Leibler model for microtubule dynamics [13] consists of two populations of microtubules undergoing dynamic instability, the longitudinal (*M*_*‖*_) and transverse population (*M*_⊥_) respectively.

#### Tubulin redistribution

The two populations compete for a finite tubulin pool, i.e., they can only access a finite number of tubulin-dimers to fuel their growth or de-novo nucleation. Let *L*_*tot*_ be the total amount of tubulin in the system, expressed as the maximum total length of microtubules to which it could give rise. Then, *L*_*tot*_ is divided over three different populations: the free tubulin pool *L*_*f*_, the tubulin incorporated into longitudinal microtubules *L*_*‖*_, and the tubulin incorporated into transverse microtubules *L*_⊥_, such that

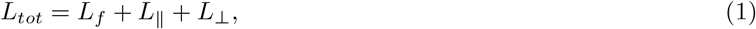

see Figure 1B. The finiteness of the tubulin pool has immediate consequences for the dynamics of microtubules. In particular, the abundance of free tubulin is positively correlated to both the nucleation rate and the growing speed of microtubules.

#### Nucleation of new microtubules

Experiments [14] have identified a Hill-type dose-response relation between the nucleation rate of new microtubules and the abundance of free tubulin, i.e.,

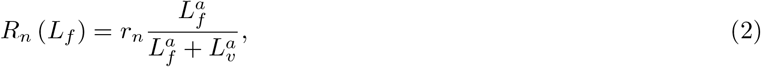

where *a* ≃ 6, *L*_*υ*_ is a constant of dimension length that governs the cross-over from a diffusion limited regime at low free tubulin densities to an intrinsic association rate limited regime at high free tubulin densities, and *r*_*n*_ is the nucleation rate of new microtubules in case of unbounded availability of tubulin.

Consistently with *in vivo* observations [15], we assume that microtubules are mainly - but not exclusively, created by microtubule-based nucleation. In particular, we divide the nucleation of new microtubules in two distinct nucleation types: microtubule-based nucleation and nucleation from a dispersed nucleation sites, with nucleation rates, respectively,

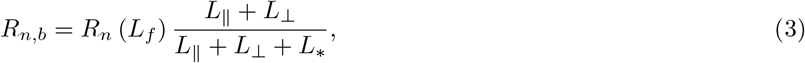

and

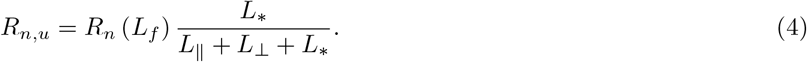

*L*_*_ is the *propensity length for dispersed nucleation*, i.e., a constant that controls the fraction of nucleation events that occur from dispersed sites in the cytosol rather than from the lattice of already existing microtubules, and a proxy for the binding affinity of nucleation complexes to the microtubule lattice.

In the CA, microtubules that arise through microtubule-based nucleation are nucleated preferentially parallel or with an angle of about 40^°^ with respect to the growth direction of the mother microtubule [3]. This nucleation mechanism by itself can already contribute to maintaining the orientation of the CA [16, 17]. As here we only consider two possible directions for microtubules, we assume that new microtubules generated through microtubule-based nucleation have a strong bias towards growing in the same direction as the mother microtubule. We include this mechanism in our model by introducing the probability *z >* 1*/*2 for a new microtubule to be nucleated parallel to the mother microtubule and, consequently, 1 − *z* to be nucleated orthogonal to it. Then, the microtubule-based nucleation rates for new longitudinal and transverse microtubules are, respectively,

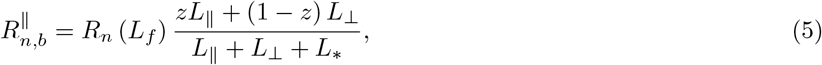

and

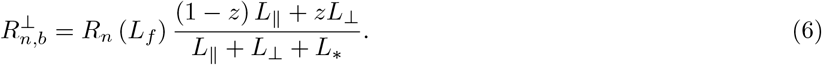

We reasonably assume that the medium is isotropic as regards the dispersed nucleation of new microtubules. Hence, the rates for dispersed nucleation are

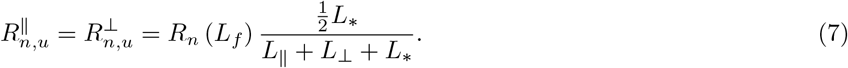

It follows that the overall nucleation rates for the two populations are 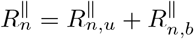 and 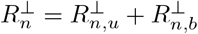 (see also Figure 1C).

#### Dynamics of individual microtubules

All microtubules are nucleated in the growing state, with growth speed *V* ^+^. Experimentally, the growth speed is approximately proportional to the amount of free tubulin [14]. Here, however, consistently with Eq. (2), we also allow for the inevitable saturation of the growth speed of microtubules, assuming it to be given by

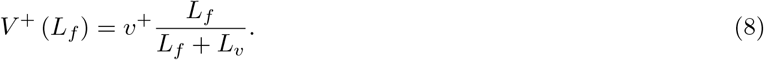

We also note that in the limit of *L*_*v*_≫1, Eq. (8) implies a linear relation between growth speed and abundance of free tubulin, in which case our model recapitulates the experimental observations. Growing microtubules can switch from the growing to the shrinking state with constant catastrophe rate *r*_*c*_. Microtubules in the shrinking state shrink with constant speed *v*^−^, and they can switch from the shrinking to the growing state with rescue rate *r*_*r*_.

When differently oriented microtubules cross each other, they create a *crossover*, where a severing event can take place. In particular, the occurrence or not of a severing event is partly influenced by the number of crossovers a microtubule has created and, consequently, to its length, and also partly by its relative position to the crossing microtubule. Indeed, experiments have shown that newer microtubules, that cross over the top of already existing ones, are preferentially severed [4]. Since our model consists of two populations of microtubules without a specific localization in space, we introduce an effective severing rate for an individual microtubule to model the creation of crossovers and the occurrence of severing events, which is proportional to the product of its own length and the total lengths of microtubules in the direction perpendicular to it. Furthermore, to be consistent with the experimental observations that new microtubules are, in general, longitudinally oriented, we introduce a preferential severing for the latter by assuming a probability 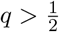 of severing a longitudinal microtubule given that a severing event has occurred. Then, if *r*_*s*_ is the intrinsic severing rate at a crossover, the overall severing rates for longitudinal and transverse microtubules of length *l* are, respectively,

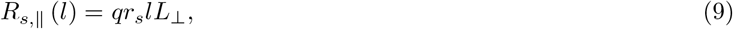

and

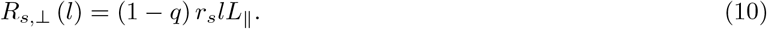

Once a microtubule is severed, the newly-created plus end of the lagging microtubule enters either the growing state with probability of stabilization-after-severing *p*^+^, or the shrinking state with probability 1 − *p*^+^, while the state of the plus end of the leading microtubule is unaffected. Experimental and theoretical work has shown that a relatively high probability of stabilization-after-severing (*p*^+^ ≃ 0.15) is required in order to commence the reorientation process [6, 7]. Finally, the newly-created minus end of the leading microtubule remains stable, without undergoing dynamic instability, see Figure 1D.

### Small increase in the growth speed of longitudinal microtubules

Previous experimental work on the effect of cell geometry on microtubule dynamics have revealed that the mechanical stress induced by cell wall geometry can influence the alignment of cortical microtubules [8, 9]. Furthermore, several computational approaches have shown that slight directional cues caused by, e.g., catastrophes or microtubule stabilization induced by features of the surface geometry, can select a single preferential alignment of the CA [11, 12].

Generically, plant cells have a cylindrical morphology. This implies that they have a clearly distinguished direction of maximal wall curvature, transverse to the cell axis, and of minimal curvature, parallel to the cell axis. It is conceivable that this small but significant difference has an impact on microtubule dynamics. We take this observation as the starting point for our main hypothesis assuming that longitudinal microtubules grow slightly faster than transverse microtubules, and that such a bias is a sufficient ingredient to obtain a maintained reorientation of the CA. Therefore, we set

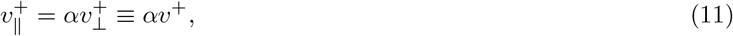

with *α >* 1 and *α ≃* 1. All other dynamic parameters of the model are unaffected. Our choice of applying a difference only in the growth speed is motivated by a practical reason. Indeed, changing only the growth speed makes the model also analytically tractable. Nevertheless, in the Results section we will show that decreasing the catastrophe rate in the longitudinal direction, which has an analogous effect on the relative stability of the transverse and longitudinal microtubules, leads to qualitatively similar results on the steady-state properties, suggesting that the choice of the specific dynamic parameter to vary in order to obtain a maintained reorientation is arbitrary.

### Polarization and transverse suppression

We refer to the initial number of transverse microtubules as 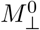, and to the amount of tubulin incorporated in them as 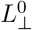. To evaluate the efficiency of the reorientation, we define the following order parameters:

- *number* and *length polarization*, respectively,

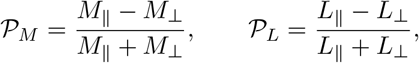

as order parameters for the difference between the two populations,
- *number* and *length transverse suppression*, respectively,

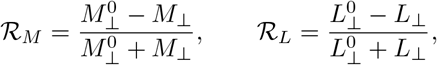

as order parameters that estimate how much of the original transverse array is still present at the end of the process.

Ideally, to consider the reorientation as efficient, we have two requirements: (i) that all four order parameters are close to 1, i.e., the majority of microtubules and the amount of incorporated tubulin are polarized in the longitudinal direction, and (ii) that the time scale for the array to reorient is comparable to the experimentally observed one [4, 6].

## III. RESULTS

### A. Computational approach

We run stochastic simulations of our model to assess whether the difference in growth speed between the two different populations is sufficient to achieve a full and maintained reorientation of the array. Simulations consist of an initial phase in which we obtain a steady-state transverse array, see Supplementary Information IX B. Then, at time *t* = 0, we allow the nucleation of longitudinal microtubules both through microtubule-based nucleation and dispersed nucleation. We let the simulations run until the system reaches again a steady-state and record the number and length polarization and suppression, and the time required to obtain a full reorientation of the CA. We perform a sensitivity analysis in the (*q, p*^+^) plane by separately tuning them from 0 to 1. We average the results over *N* = 10^3^ stochastic simulations *per* (*q, p*^+^) couple. Parameters and relative numerical values used in the simulations are listed in Table I.

**Table I.**
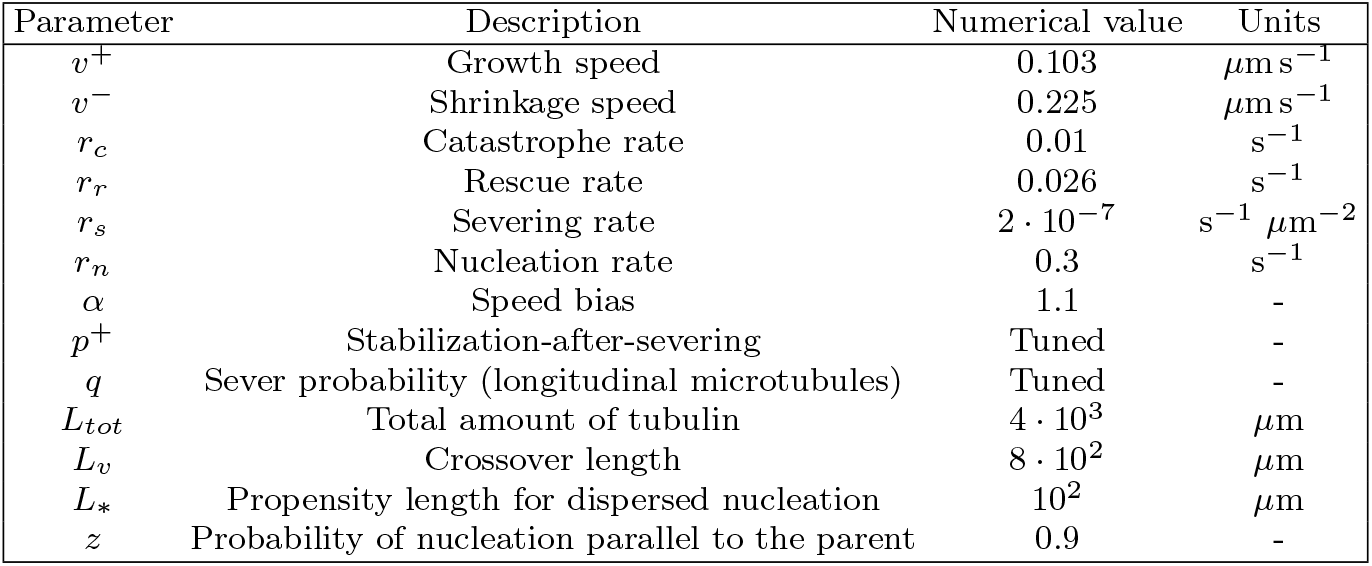
Reference values for the parameters of the model. Numerical values for the dynamic parameters and the amount of tubulin has been chosen consistently with experimental measurements [6, 14].

| Parameter | Description | Numerical value | Units |
| --- | --- | --- | --- |
| $v^+$ | Growth speed | 0.103 | $\mu\text{m s}^{-1}$ |
| $v^-$ | Shrinkage speed | 0.225 | $\mu\text{m s}^{-1}$ |
| $r_c$ | Catastrophe rate | 0.01 | $\text{s}^{-1}$ |
| $r_r$ | Rescue rate | 0.026 | $\text{s}^{-1}$ |
| $r_s$ | Severing rate | $2 \cdot 10^{-7}$ | $\text{s}^{-1} \mu\text{m}^{-2}$ |
| $r_n$ | Nucleation rate | 0.3 | $\text{s}^{-1}$ |
| $\alpha$ | Speed bias | 1.1 | - |
| $p^+$ | Stabilization-after-severing | Tuned | - |
| $q$ | Sever probability (longitudinal microtubules) | Tuned | - |
| $L_{tot}$ | Total amount of tubulin | $4 \cdot 10^3$ | $\mu\text{m}$ |
| $L_v$ | Crossover length | $8 \cdot 10^2$ | $\mu\text{m}$ |
| $L_*$ | Propensity length for dispersed nucleation | $10^2$ | $\mu\text{m}$ |
| $z$ | Probability of nucleation parallel to the parent | 0.9 | - |

Figure 2ABCD shows the polarization and the transverse suppression for both microtubule number and length. Although the figure shows that a more efficient and fast reorientation requires high values of both the parameters governing preferential severing (*q*) and stabilization-after-severing (*p*^+^), we can still observe a good degree of reorientation for *p*^+^ in the range from 0 to 0.25, i.e., in the biologically relevant range of values for the probability of stabilization-after-severing. In particular, the redistribution of tubulin seems to be very efficient in that range of values, see Figure 2BD. It is also interesting to notice that in this regime, polarization and suppression seem to be not strongly dependent on *q*. This suggests that, even though the preferential severing plays an important role in the amplification phase of the reorientation to boost the creation of longitudinal microtubules, in presence of a growth-speed bias it is not strictly required to maintain the new longitudinal array.

**Figure 2.**
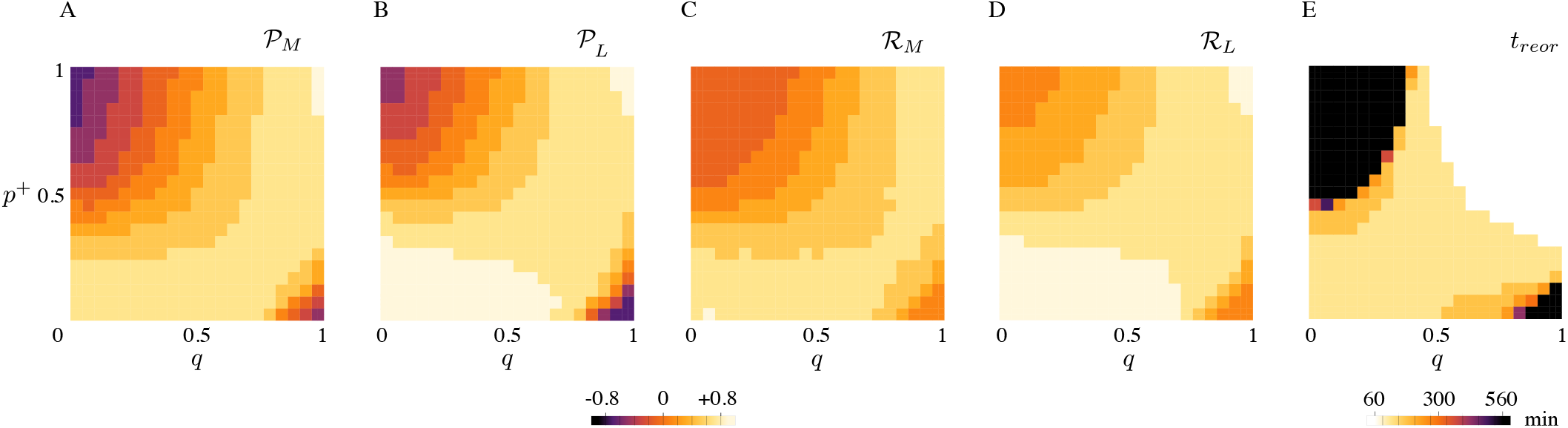
(A) Microtubule number polarization, (B) microtubule length polarization, (C) transverse number suppression, (D) transverse length suppression, (E) and transverse-to-longitudinal reorientation time as functions of *q* and *p*^+^. Lighter colors correspond to a more efficient reorientation. (A, B, C, D) The range of values for polarization and suppression runs from −1 to+1. (E) Black areas in the (*q, p*^+^) plane correspond both to reorientation processes that required more than 540 minutes, or non-occurred reorientation. Results are averaged over *N* = 10^3^ simulations.

Figure 3 shows the time evolution of the number and the total length of microtubules belonging to the different populations. The combination of preferential severing and biased growth speed has a double effect: it makes the reorientation efficient by increasing the number of longitudinal microtubules at the steady-state and suppressing the transverse, and it boosts the speed at which the reorientation occurs. Moreover, Figure 3B also shows that there is no appreciable difference between the amount of free tubulin at the beginning and at the end of the process. This means that the building material used by the longitudinal array comes from the initial transverse array rather than from the pool of free tubulin.

**Figure 3.**
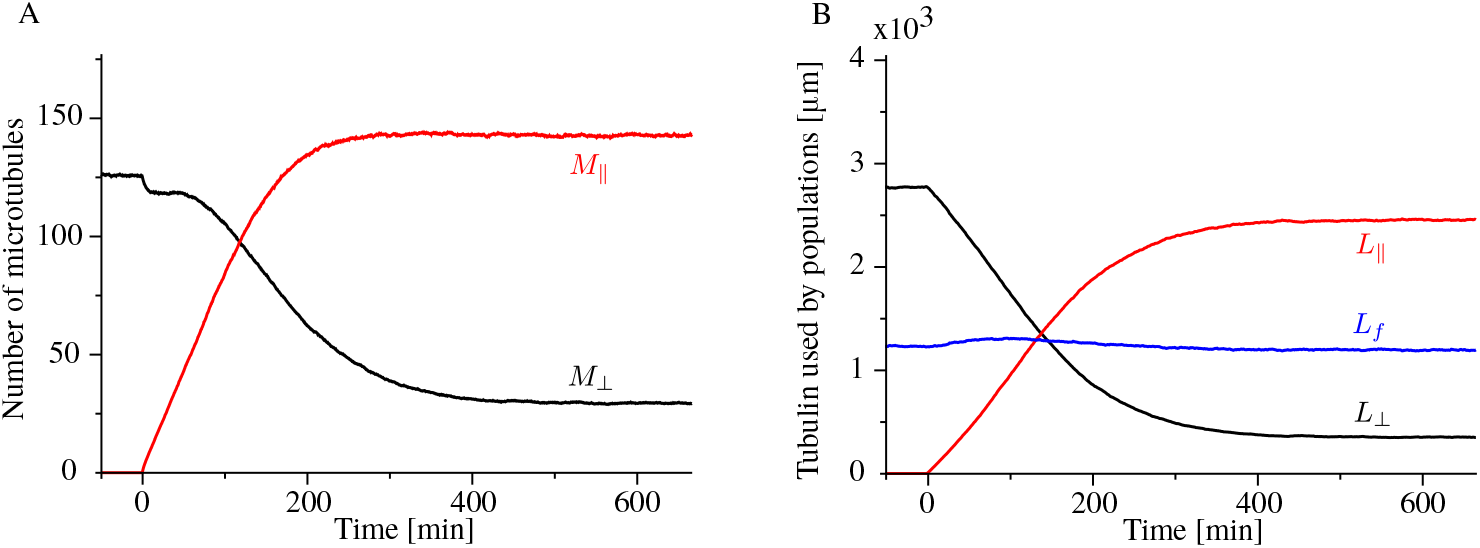
Time evolution of (A) longitudinal (red) and transverse (black) microtubules, and (B) tubulin used by the longitudinal population (red), the transverse population (black), and the free tubulin (blue), for *q* = 0.75, *p*^+^ = 0.15. Results are averaged over *N* = 10^3^ simulations.

We also tested the alternative hypothesis that the factors that promote the commencement of the reorientation process, namely, high probability of stabilization-after-severing and preferential severing for longitudinal microtubules [6, 7], could be sufficient to explain the experimentally observed maintained reorientation. Our simulations showed that, although the two ingredients are indeed necessary to quickly start the reorientation process and reach a steady-state, they cannot explain how the full reorientation is achieved and maintained (see Supplementary Information IX C).

### B. Analytical approach

To further test our hypothesis that reorientation and maintenance of the array are caused by a biased recruitment of tubulin towards the longitudinal microtubules, we analytically study a simplified version of our model in which the two populations of microtubules compete for the tubulin pool without direct interaction between them, i.e., without possibility of severing events. Therefore we set *r*_*s*_ = 0. We make the further simplification of assuming complete depolymerization of microtubules just after catastrophe events, i.e.,

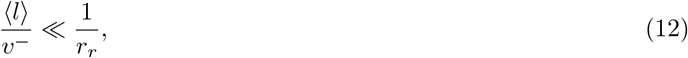

where ⟨ *l* ⟩ is the average length of microtubules. Finally, we assume that all microtubules are nucleated in the same direction of the mother microtubule, i.e., *z* = 1. To ease the notation, we hereafter drop the direct dependency of *V* ^+^ and *R*_*n*_ on *L*_*f*_ whenever this is not strictly necessary,. Under these assumptions we can rewrite the dynamic Eqs. (37-40) as

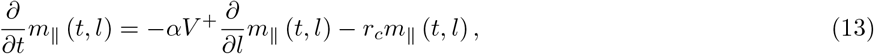

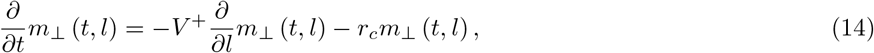

with boundary conditions

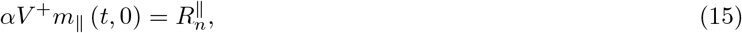

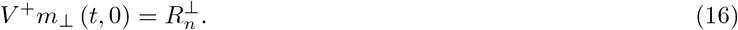

We now consider the 0^th^ and the 1^st^ moment equations corresponding to Eqs. (13) and (14), i.e.

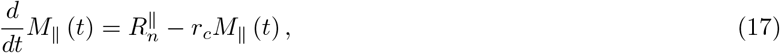

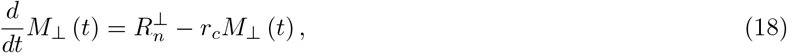

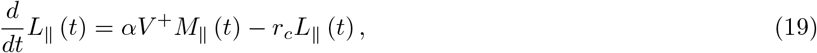

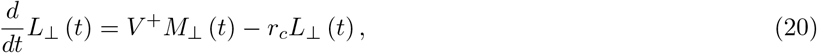

coupled with the conservation of total tubulin

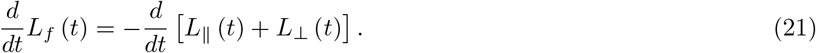

Suppose that *α* = 1. Then, becaulabelanalytical-approachse of the symmetry between longitudinal and transverse microtubules, when longitudinal microtubules start to be nucleated the overall nucleation rate of all microtubules, as well as their growth rates, remain the same as in the initial system with only the transverse array. Therefore we can safely assume that, in that specific case, *L*_*f*_ = const. Here, since *α* is slightly greater than 1, and making the reasonable assumption that *L*_*f*_ is smooth in *α*, it follows that

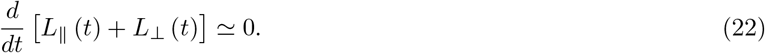

This last equation implies that all building material used by the newer longitudinal array comes from the already existing transverse one, in agreement with our computational findings, see Figure 3 B.

In order to determine what controls the ultimate polarization of the microtubule distribution, and hence the reorientation mechanism, we study the steady-state version of the moment Eqs. (17-20). For the time-dependent solution of the model, see Supplementary Information IX D. If we isolate *r*_*c*_*M* _‖_ and *r*_*c*_*M*_⊥_ from the four equations we obtain

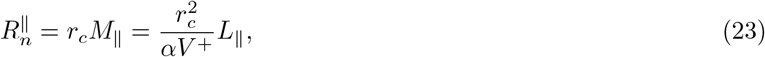

and

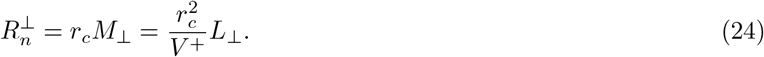

From Eq. (23) we can observe that multiplying *v*^+^ by *α* for the longitudinal growing speed is equivalent to dividing *r*_*c*_ by 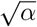.

If we define

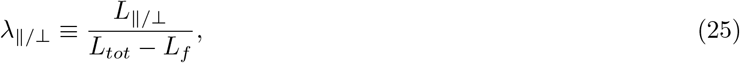

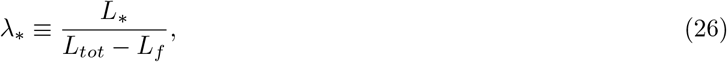

and we divide Eq. (23) by (24), by making use of Eqs. (5), (6), and (7) we obtain the system

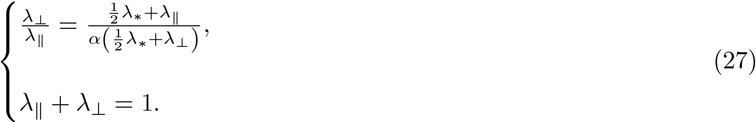

The system can be solved to find

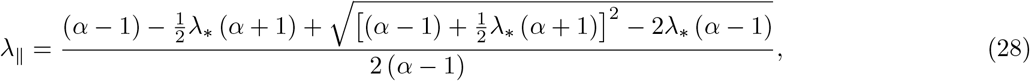

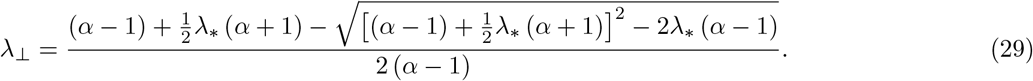

If we divide both sides of Eq. (29) by 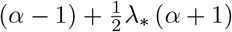 and we expand the square root we obtain

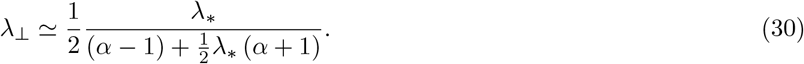

Consequently

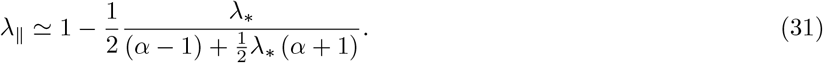

Inserting Eqs. (30) and (31) into Eqs. (23) and (24), we obtain the number of microtubules in the longitudinal and in the transverse directions

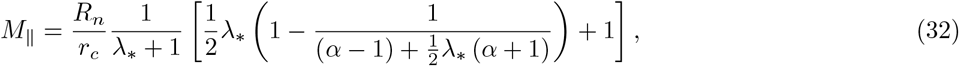

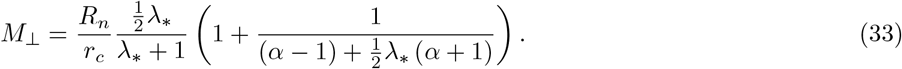

Notice that, in the *λ*_*_ → 0 limit - i.e., when nucleation is only microtubule-based, all tubulin is polarized in the longitudinal direction, and we observe full reorientation of the array from the transverse to the longitudinal direction. Indeed 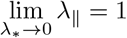, and 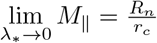. On the other hand, when *λ*_*_ → *∞* - i.e., when nucleation is only dispersed,

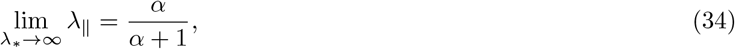

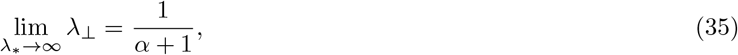

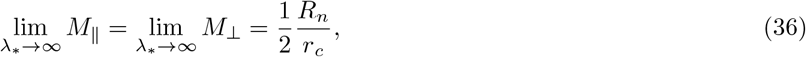

implying that, although in this limit the isotropy of the dispersed nucleation imposes that the final number of microtubules of the two populations is the same, the *α >* 1 bias in the growth speed still produces a slight tubulin polarization in the longitudinal direction by virtue of the longitudinal microtubules being on average slightly longer than the transverse ones, see Figure 4 and Supplementary Information IX E. The foregoing analysis shows that even in the absence of the severing process, the bias in the growth speed by itself is a sufficient driver of the shift towards the longitudinally polarized state.

**Figure 4.**
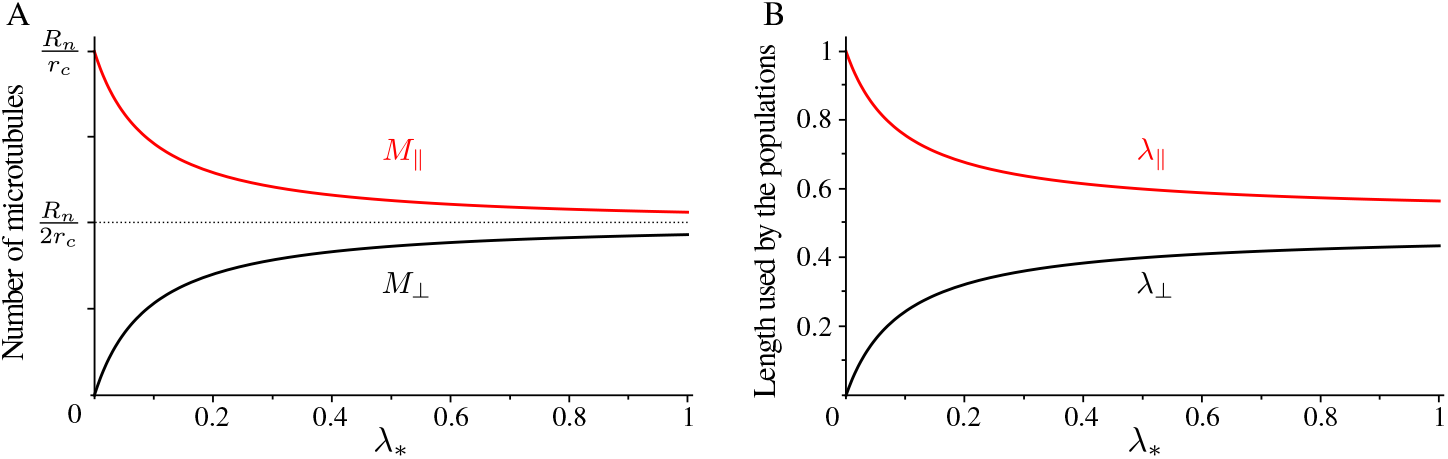
(A) Number of microtubules and (B) non-dimensional length used by microtubules populations as functions of the propensity length for dispersed nucleation, with *α* = 1.1.

## IV. DISCUSSION

Our goal, here, was to propose a plausible mechanism by which the striking reorientation of the CA of dark-grown hypocotyl cells upon exposure to blue light can be “locked-in” on longer post-stimulus timescales. A major ingredient of the model we propose, the fact that tubulin availability is an important limiting factor, follows naturally from the mechanism originally put forward to explain the onset reorientation, which involves the exponential amplification of longitudinal microtubules due to the initially predominant severing occurring at cross-overs with the pre-existing transverse array. This exponential increase will, at some time of necessity, deplete the pool of available tubulin. From that point on, the process essentially chokes itself and an inevitable redistribution of orientations towards a more isotropic equilibrium distribution follows. This points to the need for an additional assumption that explains the ultimate dominance of the longitudinal microtubules, which we provide in the form of the hypothesis that longitudinal microtubules are dynamically advantaged, be it through a slightly higher base growth speed, or a slightly decreased intrinsic catastrophe rate. This makes our proposed mechanism an interesting example of the generic competitive exclusion principle in population ecology [18]. According to that principle, two populations competing for the same limited resources cannot coexist at equilibrium. Instead, the competitively superior population will persist, while the other will go extinct. In our case, the strict validity of this principle is attenuated by three regulating factors that prevent the total extinction of the initial transverse array: the occasional occurrence of severing events for transverse microtubules, the few dispersed nucleation events in the transverse direction, and the microtubule-based nucleation of transverse from longitudinal microtubules. However, when none of those three factors is present, as in the *λ*_*_ → 0 case of Section III B, only the longitudinal array would persist.

It is somewhat striking that preferential severing for longitudinal microtubules and high probability of stabilization-after-severing, two factors which have earlier been identified as crucial for the start of the reorientation process [6, 7], do not appear to be necessary for the long-term maintenance of the longitudinal array. However, the computational results presented here reveal that both preferential severing and high probability of stabilization-after-severing play a key role in determining the speed of the reorientation.

Naturally, our proposed model raises some questions and problems, which we hope can be addressed by further research. Firstly, an experimental observation of a difference in any of the dynamic parameters between differently oriented microtubules is, at present, lacking. Nevertheless, such a difference could feasibly be revealed by a more detailed data analysis on microscopy movies of cortical microtubules of *Arabidopsis thaliana* during the reorientation of the cortical array. The detection of an even small difference between the dynamics of longitudinal and transverse microtubules could potentially provide support for our theoretical predictions. Of course, we may already speculate on possible sources of such a difference. One of these could be differences in mechanical stress to which microtubules are subject as a consequence of the cell geometry. Indeed, in a roughly cylindrical cell morphology the wall stress in the curved transverse direction is larger than in the flat longitudinal directions. Intriguingly, and at first sight at odds with our assumptions, this difference is, in fact, the basis for a currently widely accepted explanation for the predominance of the transverse direction of cortical microtubules, based on observations that microtubules consistently orient themselves along lines of maximal stress [8, 10, 19]. The molecular mechanism responsible for this preference, however, is not as yet fully elucidated. At the same time, it does point to the potential for mechanisms influencing microtubule stability that couple to the curvature of the cortex, leaving open the question whether these are enhancing or depressing depending on the magnitude of the curvature. Secondly, the model as presented here is still highly idealized, and arguably needs to be developed further to include a more realistic description of the actual system. Specifically, our assumption that the transverse and longitudinal microtubule populations effectively live in distinct “spaces” and only interact at an ensemble level through the shared tubulin pool and the mutual modulation of the severing rates is a strong simplification. In reality, the common cell geometry which they share, the fact that they have continuous orientation, as well as the full repertoire of interactions at the level of individual microtubules such as, e.g., induced catastrophes due to collisions and zippering should be taken into account. Whilst the complexity of adding this level of detail will preclude an analytical approach, the computational tools for such a more comprehensive approach are available, e.g., in the form of the framework described in [20, 21] and recently extended to deal with arbitrary cell geometries [11]. Addition of a bias in the speed of growing microtubules depending on their orientation is readily implementable and, in principle, would allow for a more realistic computational test whether such a difference can indeed explain a reorientation of the array and its long-term maintenance.

## V. AUTHOR’S CONTRIBUTION

M.S. designed and performed the simulations, and carried out the formal analysis. M.S. and B.M.M. conceived the study, designed the model, and wrote the manuscript

## VI. ACKNOWLEDGEMENT

The work of M.S. was supported by the ERC 2013 Synergy Grant MODELCELL. The work of B.M.M. is part of the research program of the Dutch Research Council (NWO). We thank Jelmer Lindeboom for a careful reading of the manuscript and for providing Figure 1A.

## VII. DATA AVAILABILITY

The simulations code that support the findings of this study is available from the corresponding author, Marco Saltini, upon reasonable request.

## VIII. CONFLICTS OF INTEREST

The authors declare no competing interests.

## IX. SUPPLEMENTARY INFORMATION

### A. Steady-state solution for the microtubule length distribution

#### Dynamic equations

Let *m*^*τ*^ (*t, l*) be the probability distribution of the length of microtubules in the state *τ* at time *t*. Following Tindemans and Mulder [22], it is possible to show that the dynamic equations that govern the model are

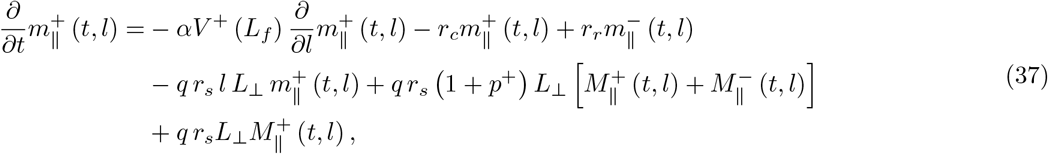

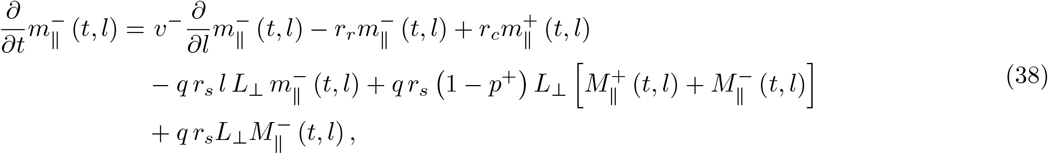

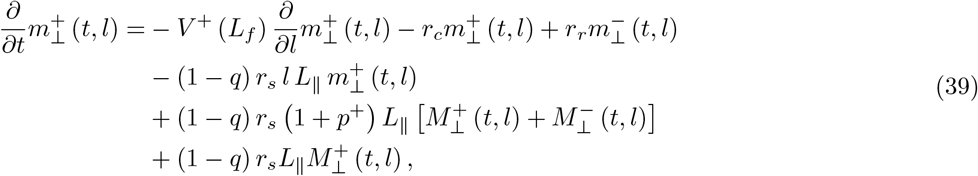

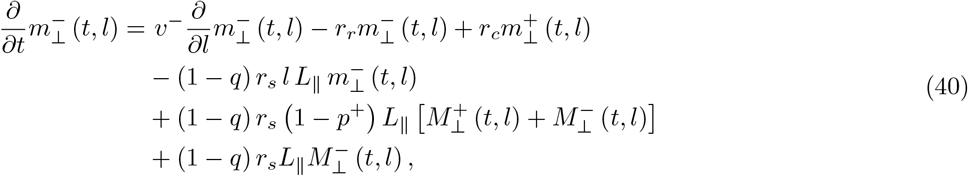

with boundary conditions

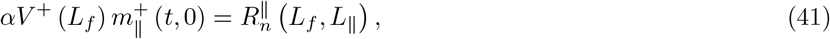

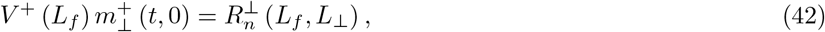

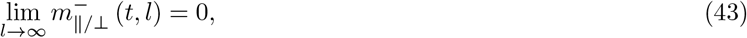

and initial conditions from the steady-state solution of the Dogterom-Leibler model, i.e.,

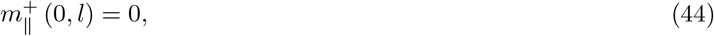

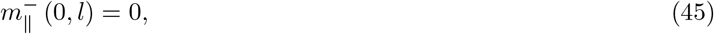

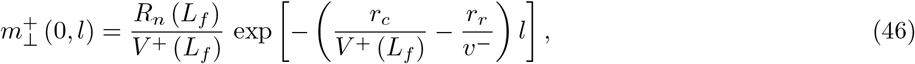

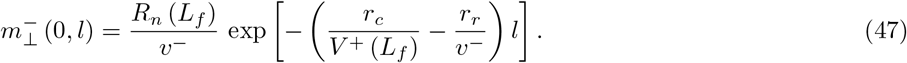

It is convenient to underline that the total amount of tubulin polarized in either of the two directions is linked to the microtubule length distributions through the equations

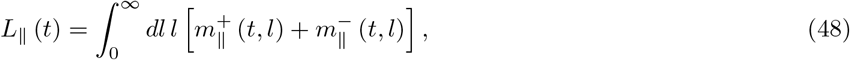

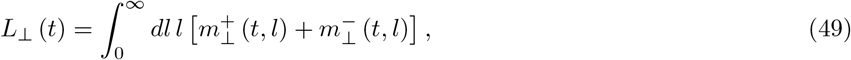

i.e., the total amount of tubulin used by longitudinal/transverse microtubules is the first moment of the total length distribution of longitudinal/transverse microtubules.

#### Steady-state solution

The dependency of the growing speed of microtubules on the amount of free tubulin *L*_*f*_ in the pool implies that the microtubule length distribution eventually reaches the steady-state. Here, in order to find an analytical solution, we make the further assumption that microtubules undergo complete depolymerization suddenly after a catastrophe, i.e.,

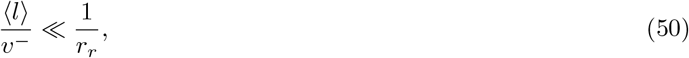

where ⟨*l* ⟩ is the mean length of a microtubule in the system. In this limit we can identify all microtubules with the growing microtubules and, therefore, to ease the notation we remove the label + from all microtubule distributions. Thus, Eqs. (37-40) are replaced by

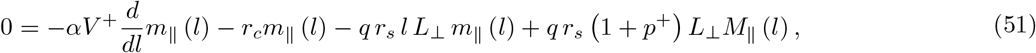

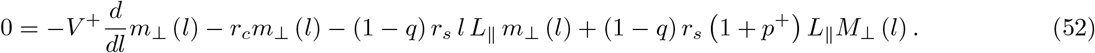

If we differentiate by *l* the last two equations we obtain

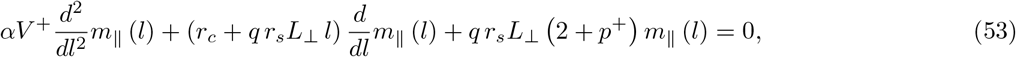

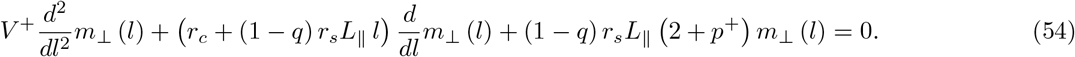

As 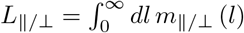, Eqs. (53) and (54) are coupled second order differential equations, and they are also linked to Eq. (1) through *V* ^+^ (*L*_*f*_). However, *L*_*f*_, *L*_*‖*_, and *L*_⊥_ do not depend on *l*, hence Eqs. (53) and (54) can be in principle solved, with solutions that depend on the total amount of tubulin used by the other population and on the free tubulin. We re-write the two equations in a more elegant way as

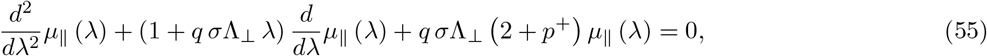

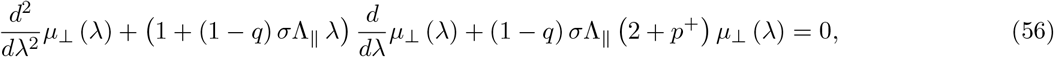

where we introduce the non-dimensional quantities

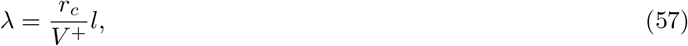

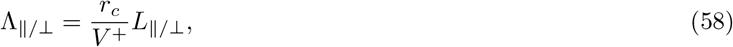

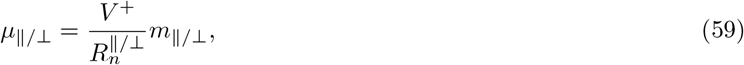

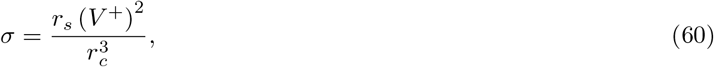

and we incorporate the factor *α* in the parameter *V* ^+^ for longitudinal microtubules. Notice that, with this non-dimensionalization, all parameters of the model - including the independent variable *λ*, are functions of *L*_*f*_ as they depend on *V* ^+^ (*L*_*f*_).

To solve Eq. (55), we first notice that the asymptotic solution for large *λ* decays as exp 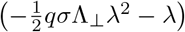. Therefore, we suppose there exists a function *ξ* _*‖*_ (*λ*) such that

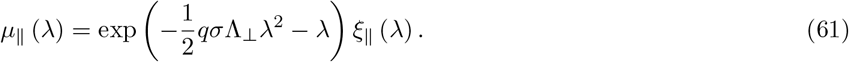

If we plug this in Eq. (55) we obtain

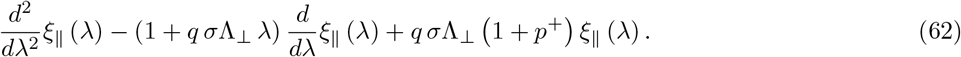

If we change variable as 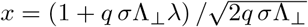, last equation becomes

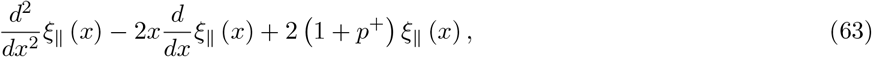

i.e., the Hermite equation [23], the solution of which is the Hermite function

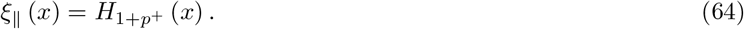

Therefore, the full solution for *µ*_*‖*_ becomes

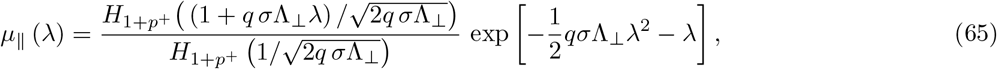

and, similarly,

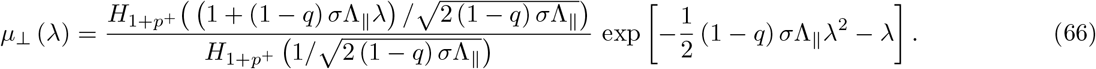

Eqs. (65) and (66) highlight a peculiar property of the model. Indeed, if *p*^+^ ≠ 0, both denominators of the two distribution can be identically 0. In other words, there exist two values of *σ* such that 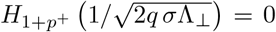 or 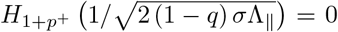, with the consequence that the number of longitudinal/transverse microtubules diverges. In the analytically tractable *p*^+^ = 1 case these values are 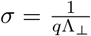 for the divergence of the longitudinal, and 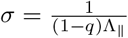 for the divergence of the transverse microtubules.

### B. The initial transverse array

To keep with the assumption coming from the experiments that initially all cortical microtubules are directed transversely to the growth direction of the cell, we build the initial array by considering dispersed and microtubule-based nucleation possible only in the transverse direction. In other words, for the creation of the initial array, we impose

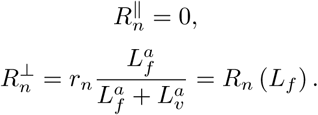

Therefore, the dynamics of the initial array is fully described by the Dogterom-Leibler model. Furthermore, the dependency of the growing speed of microtubules on the amount of free tubulin *L*_*f*_ in the pool implies that the microtubule length distribution eventually reaches the steady-state even in the case of initially unbounded-growth microtubules (Tindemans et al., Front. Phys., 2014). Thus, the solution of the model is the steady-state solution of Dogterom-Leibler model. Such a solution for the distribution of the length of microtubules is the initial condition for our model, see Supplementary Information IX A.

### C. Polarization and suppression in case of no difference in the growing speed of different populations

Here, we show that the sole asymmetry in the system consequent to the preferential severing cannot explain the full, maintained reorientation observed in the experiments. Similarly as in Result section, we perform a sensitivity analysis by separately tuning *q* and *p*^+^ from 0 to 1, and we measure number and length polarization and suppression, and the time needed by the system to achieve the reorientation, averaged over *N* = 10^3^ simulations. Parameters and relative numerical values used in the simulations are listed in Table I. However, since we are considering the case in which differently oriented microtubules have the same growing speed, in this case *α* = 1.

Figure 5ABE shows that the system does not exhibit longitudinal polarization at the equilibrium for biologically realistic *p*^+^, i.e., for *p*^+^ comprises between 0 and 0.25. Similarly, Figure 5CD shows that in the same range of values, although we can appreciate some degree of transverse suppression, the initial transverse array does not disappear. On the other hand, Figure 5 reveals the existence of two regions in the (*q, p*^+^) plane where the reorientation occurs and it is fast, namely, when both *q* and *p*^+^ are high and, surprisingly, when they are both low. While one can easily argue that a high value of both *q* and *p*^+^ is associated to a greater likelihood of increasing the size of the longitudinal population and the lifetime of their individuals and, hence, the longitudinal polarization, it is more difficult to intuitively understand the behaviour of the system for low values of those probabilities. However, lower values of *q* and *p*^+^ are linked to an effective shortening of the single transverse microtubules and, therefore, to a fall in their average lifetime. As a consequence, the overall length used by the transverse array shortens, and so does the number of transverse microtubules, as their nucleation is partly correlated to the length polarized in the transverse direction. Nevertheless, although narrow areas in the heat maps of Figure 5 where a full and maintained reorientation of the CA occurs do exist, a comparison with the Figure 2 would immediately show that in the latter case the same areas are wider.

**Figure 5.**
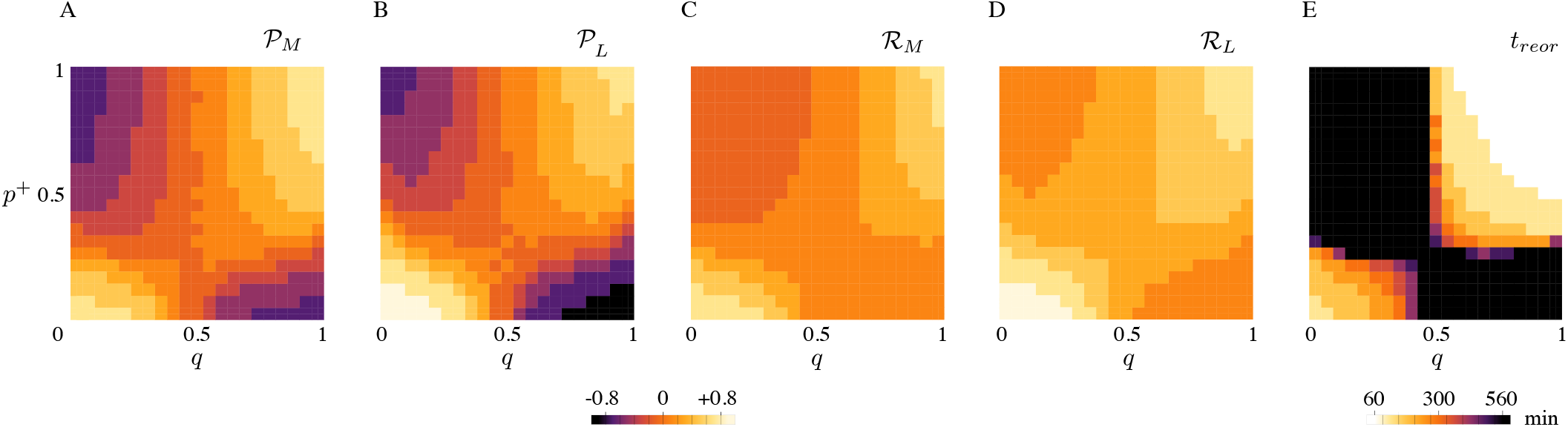
(A) Microtubule number polarization, (B) microtubule length polarization, (C) transverse number suppression, (D) transverse length suppression, (E) and transverse-to-longitudinal reorientation time as functions of *q* and *p*^+^. Lighter colors correspond to a more efficien(t reor)ientation. (A, B, C, D) The range of values for polarization and suppression runs from −1 to+1. (E) Black areas in the (*q, p*^+^) plane correspond both to reorientation processes that required more than 540 minutes, or non-occurred reorientation. Results are averaged over *N* = 10^3^ simulations.

As the only asymmetry between the two populations of the system is due to the probability *q*, Figure 5AB displays, as expected, the symmetry 𝒫 → − 𝒫, as *q* → 1 − *q* for both 𝒫_*M*_ and 𝒫_*L*_.

Although from these results one can conclude that the preferential severing for the longitudinal microtubules cannot explain the maintenance of the array in the longitudinal direction, at least in the biological range of values for *p*^+^, the dynamic behaviour of the two microtubule populations for high values of *q* and *p*^+^ highlights an interesting fact.

Indeed, Figure 5E shows that high values of *p*^+^ and *q* are associated with fast reorientation, showing that, even though the preferential severing for longitudinal microtubules and a high probability of stabilization-after-severing are neither necessary nor sufficient to achieve CA reorientation, they are able to accelerate this process significantly.

The amount of free tubulin in the pool does not substantially change from the initial value where only transverse microtubules are present, to the final steady-state value, see Figure 6B. This means that all the building material used by the longitudinal microtubules to create the new array comes from the suppression of the initial transverse array. Curiously, Figure 6A also shows that at the start of the reorientation process, i.e., at *t* = 0, there is a sudden little drop of the number of transverse microtubules. This may be explained by the sudden change in the nucleation rate of old microtubules, as it switches from *R*_*n*_ to *R*_*n*_*/*2 as a consequence of the imposed isotropy of the system.

**Figure 6.**
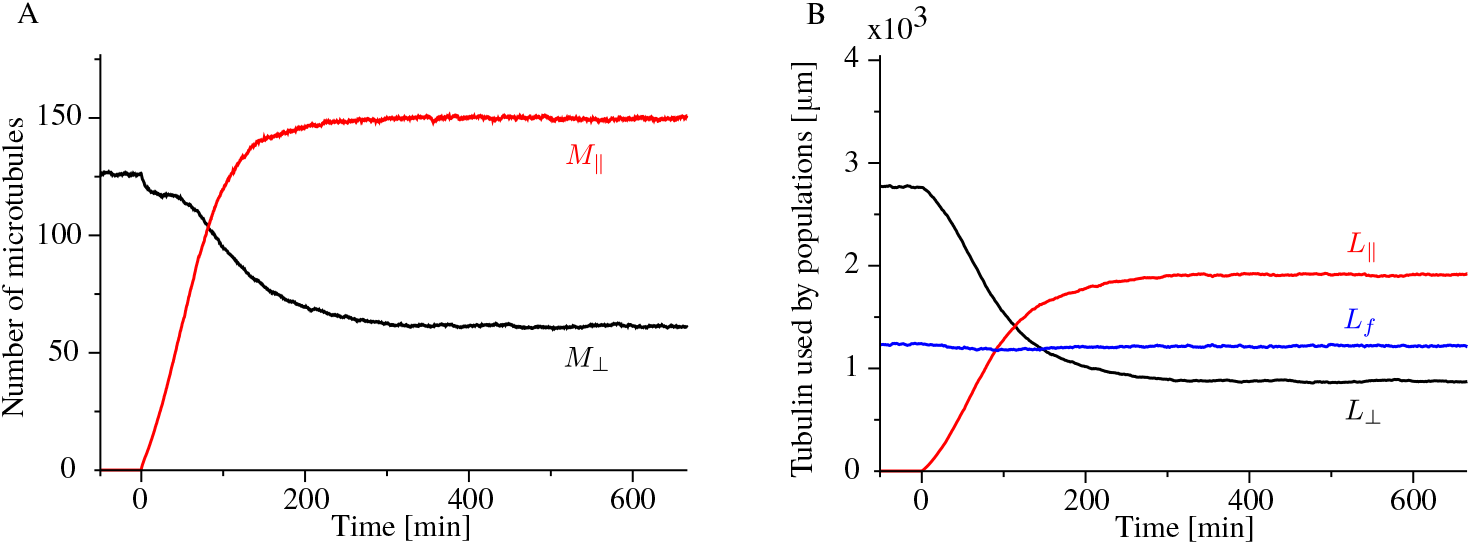
Time evolution of (A) longitudinal (red) and transverse (black) microtubules, and (B) tubulin used by the longitudinal population (red), the transverse population (black), and the free tubulin (blue), averaged over *N* = 10^3^ simulations. Here, we used *q* = 0.8 and *p*^+^ = 0.6.

### D. Time-dependent solution of the model without severing events

Here, we study the time scale of the reorientation of the array from the transverse to the longitudinal direction for the model discussed in Section III B, i.e., when no severing events can occur (*r*_*s*_ = 0) and microtubules depolymerize completely after a catastrophe (⟨ *l* ⟩ */v*^−^ ≪ 1*/r*_*r*_). We focus on Eqs. (17) and (19), with their respective boundary conditions

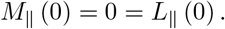

By plugging these boundary conditions in Eqs. (17) and (19) we obtain the boundary conditions for the first derivative of both *M*_‖_ and *L*_‖_

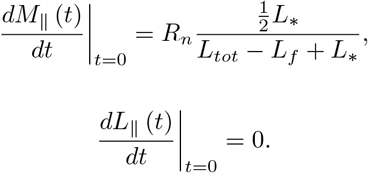

Eqs. (17) and (19) can be decoupled to obtain

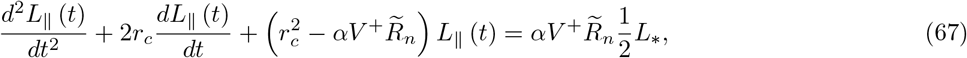

and

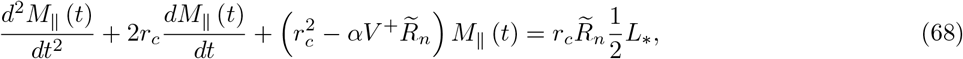

where we defined

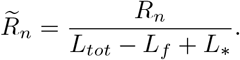

Eqs. (67) are second order non-homogeneous differential equations, the solutions of which are

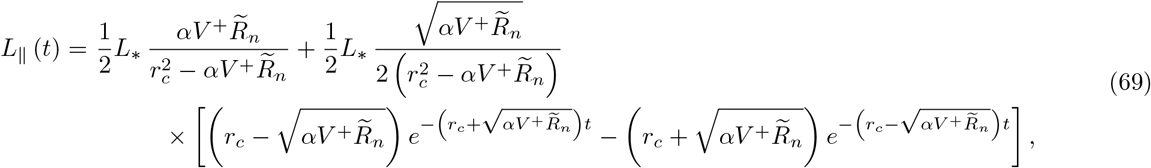

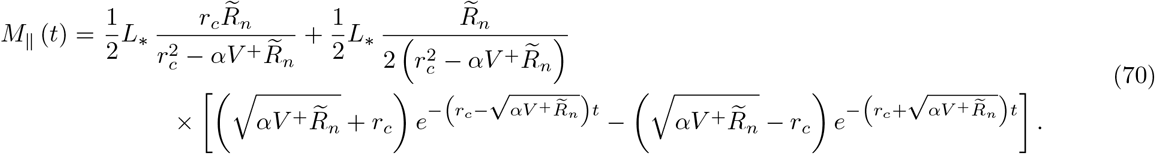

These expressions define the time scale of the reorientation process, i.e.

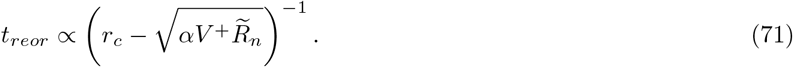

To highlight all dependencies of *t*_*reor*_ on the model parameters, we can conveniently rewrite 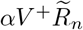 by expressing all quantities as functions of *L*_*f*_. We find

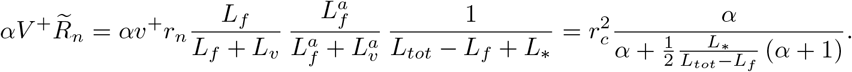

Thus, the time scale of the reorientation can be written now as

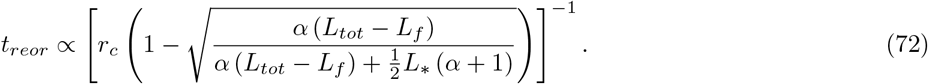

*L*_*f*_ is weakly dependent on *r*_*c*_. As a consequence, Eq. (72) reveals that the time scale of the reorientation is inversely proportional to the catastrophe rate. The interpretation of this counter-intuitive result, is that every time a microtubule undergoes a catastrophe, it releases to the free tubulin pool an amount of tubulin equal to its length. Therefore, the amount of building material available for the new array increases with higher rate, and so does the speed of reorientation.

### E. Polarization and transverse suppression in the model without severing events

We calculate the reorientation polarization and transverse suppression for the model discussed in Section III B, i.e., when no severing events can occur (*r*_*s*_ = 0) and microtubules depolymerize completely after a catastrophe (⟨*l*⟩*) /υ*^−^ ≪ 1*/r*_*r*_). From Eqs. (30-33) we obtain

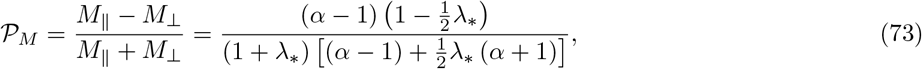

and

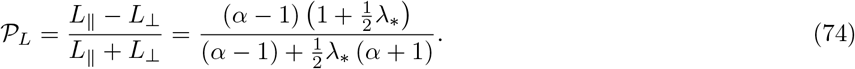

As expected, both reorientation parameters are close to 1 when *λ*_*_ is small, see Figure 7AB, whilst they rapidly decay to 0 for higher values of *λ*_*_.

**Figure 7.**
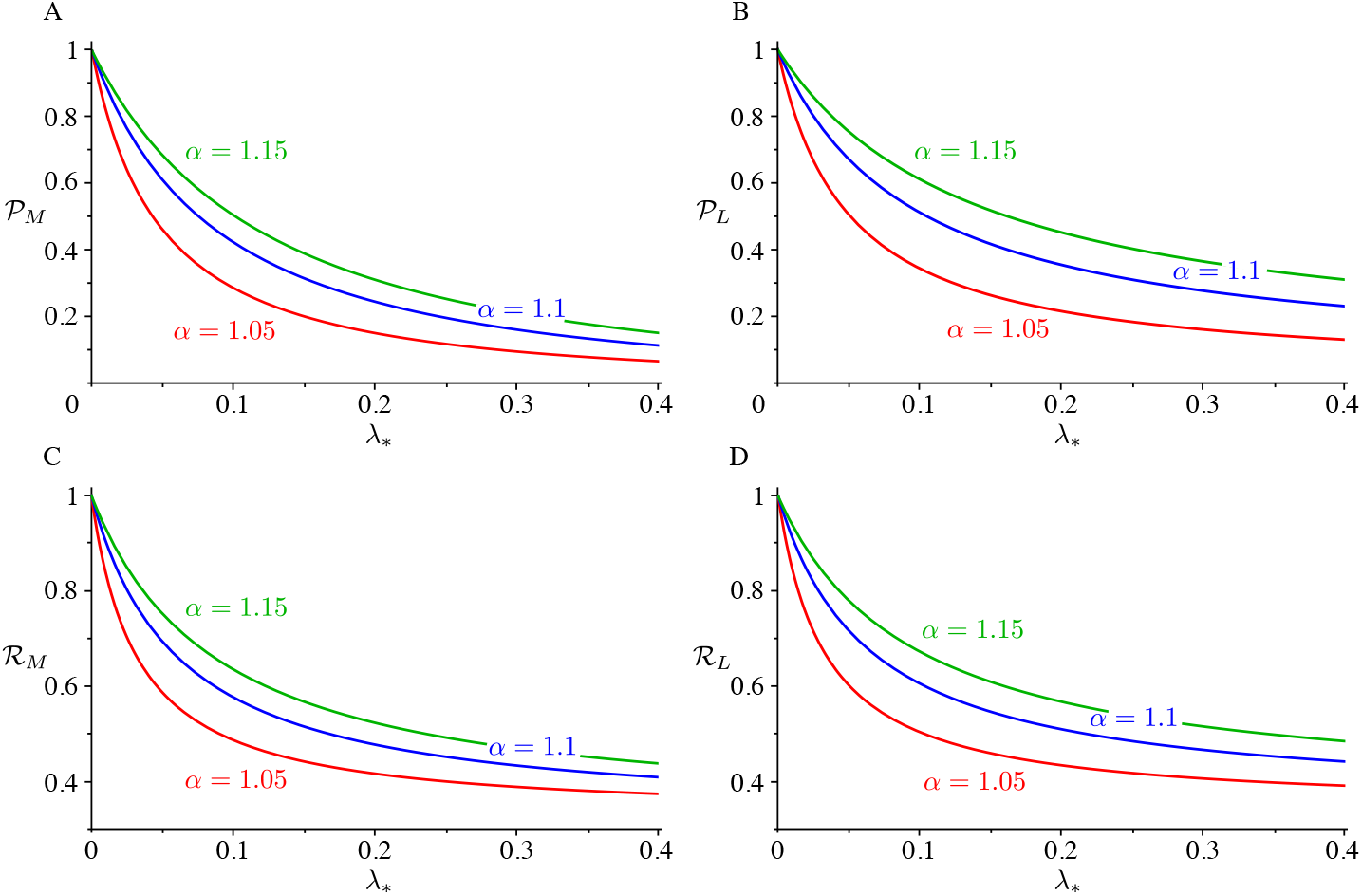
Polarization for (A) the microtubules and (B) the tubulin, and suppression for (C) microtubules number and (D) length as a function of the propensity length for dispersed nucleation for *α* = 1.05 (red), *α* = 1.1 (blue), and *α* = 1.15 (green).

Similarly, we can calculate the transverse suppression:

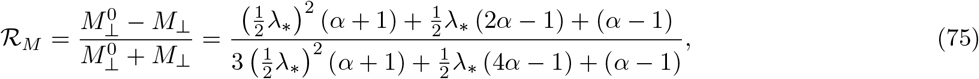

and

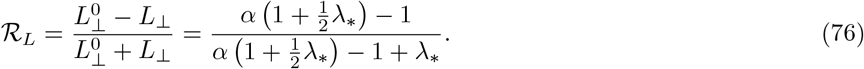

Figure 7CD shows that in the case of the transverse suppression the effect of the bias in the speed is to almost completely remove the initial transverse array.

## Notes

### Competing Interest Statement

The authors have declared no competing interest.

## References

[1] Ram Dixit and Richard Cyr. The cortical microtubule array: From dynamics to organization, oct 2004.

[2] Takashi Murata, Seiji Sonobe, Tobias I. Baskin, Susumu Hyodo, Seiichiro Hasezawa, Toshiyuki Nagata, Tetsuya Horio, and Mitsuyasu Hasebe. Microtubule-dependent microtubule nucleation based on recruitment of γ-tubulin in higher plants. Nat. Cell Biol., 7(10):961–968, oct 2005.

[3] Jordi Chan, Adrian Sambade, Grant Calder, and Clive Lloyd. Arabidopsis cortical microtubules are initiated along, as well as branching from, existing microtubules. Plant Cell, 21(8):2298–2306, aug 2009.

[4] Jelmer J. Lindeboom, Masayoshi Nakamura, Anneke Hibbel, Kostya Shundyak, Ryan Gutierrez, Tijs Ketelaar, Anne Mie C. Emons, Bela M. Mulder, Viktor Kirik, and David W. Ehrhardt. A mechanism for reorientation of cortical microtubule arrays driven by microtubule severing. Science (80-.)., 342(6163), ec 2013.

[5] Masayoshi Nakamura, Jelmer J. Lindeboom, Marco Saltini, Bela M. Mulder, and David W. Ehrhardt. SPR2 protects minus ends to promote severing and reorientation of plant cortical microtubule arrays. J. Cell Biol., 217(3):915–927, mar 2018.

[6] Jelmer J. Lindeboom, Masayoshi Nakamura, Marco Saltini, Anneke Hibbel, Ankit Walia, Tijs Ketelaar, Anne Mie C. Emons, John C. Sedbrook, Viktor Kirik, Bela M. Mulder, and David W. Ehrhardt. CLASP stabilization of plus ends created by severing promotes microtubule creation and reorientation. J. Cell Biol., 218(1):190–205, jan 2019.

[7] Marco Saltini and Bela M. Mulder. Critical threshold for microtubule amplification through templated severing. Phys. Rev. E, 101(5):052405, may 2020.

[8] Olivier Hamant, Marcus G. Heisler, Henrik Jönsson, Pawel Krupinski, Magalie Uyttewaal, Plamen Bokov, Francis Corson, Patrik Sahlin, Arezki Boudaoud, Elliot M. Meyerowitz, Yves Couder, and Jan Traas. Developmental patterning by mechanical signals in arabidopsis. Science, 322(5908):1650–1655, 2008.

[9] Benoît Landrein and Olivier Hamant. How mechanical stress controls microtubule behavior and morphogenesis in plants: history, experiments and revisited theories. Plant J., 75(2):324–338, jul 2013.

[10] Magalie Uyttewaal, Agata Burian, Karen Alim, Benoît Landrein, Dorota Borowska-Wykr ęt, Annick Dedieu, Alexis Peaucelle, Michal Ludynia, Jan Traas, Arezki Boudaoud, Dorota Kwiatkowska, and Olivier Hamant. Mechanical stress acts via katanin to amplify differences in growth rate between adjacent cells in arabidopsis. Cell, 149(2):439–451, 2012.

[11] Bandan Chakrabortty, Ikram Blilou, Ben Scheres, and Bela M. Mulder. A computational framework for cortical microtubule dynamics in realistically shaped plant cells. PLoS Comput. Biol., 14(2):e1005959, feb 2018.

[12] Vincent Mirabet, Pawel Krupinski, Olivier Hamant, Elliot M. Meyerowitz, Henrik Jönsson, and Arezki Boudaoud. The self-organization of plant microtubules inside the cell volume yields their cortical localization, stable alignment, and sensitivity to external cues. PLOS Computational Biology, 14(2):1–23, 02 2018.

[13] Marileen Dogterom and Stanislas Leibler. Physical aspects of the growth and regulation of microtubule structures. Phys. Rev. Lett., 70(9):1347–1350, mar 1993.

[14] Michal Wieczorek, Susanne Bechstedt, Sami Chaaban, and Gary J. Brouhard. Microtubule-associated proteins control the kinetics of microtubule nucleation. Nat. Cell Biol., 17(7):907–916, jul 2015.

[15] David W. Ehrhardt. Straighten up and fly right-microtubule dynamics and organization of non-centrosomal arrays in higher plants, feb 2008.

[16] Eva E. Deinum, Simon H. Tindemans, and Bela M. Mulder. Taking directions: The role of microtubule-bound nucleation in the self-organization of the plant cortical array. Phys. Biol., 8(5), oct 2011.

[17] Panayiotis Foteinopoulos and Bela M. Mulder. The Effect of Anisotropic Microtubule-Bound Nucleations on Ordering in the Plant Cortical Array. Bulletin of Mathematical Biology, 76(11):2907–2922, nov 2014.

[18] Garrett Hardin. The competitive exclusion principle. Science (80-.)., 131(3409):1292–1297, apr 1960.

[19] Leia Colin, Antoine Chevallier, Satoru Tsugawa, Florian Gacon, Christophe Godin, Virgile Viasnoff, Timothy E. Saunders, and Olivier Hamant. Cortical tension overrides geometrical cues to orient microtubules in confined protoplasts. Proceedings of the National Academy of Sciences, 117(51):32731–32738, 2020.

[20] Simon H. Tindemans, Rhoda J. Hawkins, and Bela M. Mulder. Survival of the aligned: Ordering of the plant cortical microtubule array. Phys. Rev. Lett., 104(5):058103, feb 2010.

[21] Simon H. Tindemans, Eva E. Deinum, Jelmer J. Lindeboom, and Bela M. Mulder. Efficient event-driven simulations shed new light on microtubule organization in the plant cortical array. Front. Phys., 2:1–15, apr 2014.

[22] Simon H. Tindemans and Bela M. Mulder. Microtubule length distributions in the presence of protein-induced severing. Phys. Rev. E - Stat. Nonlinear, Soft Matter Phys., 81(3):031910, mar 2010.

[23] Richard Courant and David Hilbert. Methods of Mathematical Physics. Wiley-Interscience, 1989.

